# Assessing Individual Sensitivity to the Thermal Grill Illusion: A Two-Dimensional Adaptive Psychophysical Approach

**DOI:** 10.1101/2024.03.14.585017

**Authors:** Camila Sardeto Deolindo, Jesper Fischer Ehmsen, Arthur S. Courtin, Alexandra G. Mitchell, Camilla E. Kraenge, Niia Nikolova, Micah G. Allen, Francesca Fardo

**Affiliations:** Center of Functionally Integrative Neuroscience (CFIN), Department of Clinical Medicine, Aarhus University; Cambridge Psychiatry, University of Cambridge, United Kingdom; Danish Pain Research Center, Department of Clinical Medicine, Aarhus University

**Keywords:** Pain, Thermosensation, Psychophysics, Thermal Grill Illusion, 2D-TGC

## Abstract

In the thermal grill illusion (TGI), the spatial alternation of non-noxious warm and cold temperatures elicits burning sensations that resemble the presence of noxious stimuli. Previous research has largely relied on the use of specific temperature values (i.e., 20°C and 40°C) to study this phenomenon in both healthy individuals and patient populations. However, this methodology fails to account for inter-individual differences in thermal sensitivity, limiting the precision with which TGI responses can be evaluated across diverse populations. To address this gap, we created a Two-Dimensional Thermal Grill Calibration (2D-TGC) protocol, enabling an efficient and precise estimation of the combinations of warm and cold temperatures needed to elicit burning sensations tailored to each individual. By applying the 2D-TGC protocol in 43 healthy participants, we demonstrated key findings: (1) The TGI can be thresholded using an adaptive psychophysical method. (2) Multiple combinations of warm and cold temperatures can elicit this phenomenon. (3) The protocol facilitated the identification of temperature combinations that elicit TGI with varying levels of probability, intensity, and perceived quality ranging from freezing cold to burning hot. (4) TGI responsivity can be quantified as a continuous variable, moving beyond the conventional classification of individuals as responders vs. non-responders based on arbitrary temperature values. The 2D-TGC offers a comprehensive approach to investigate the TGI across populations with altered thermal sensitivity, and can be integrated with other methods (e.g., neuroimaging) to elucidate the mechanisms responsible for perceptual illusions in the thermo-nociceptive system.

## Introduction

Perceptual illusions represent a category of phenomena where the subjective experience of a stimulus diverges from its physical characteristics. Illusions challenge conventional understanding of perception, illustrating how sensory processing creates percepts that are seemingly counterintuitive with respect to the actual properties of a stimulus. Far from being mere quirks or deceits of sensory processing, illusions reveal consistent and quantifiable responses, both in terms of the conditions that elicit them and their specific perceptual attributes [1].

The field of psychophysics has been pivotal in characterizing perceptual illusions, utilizing rigorous methodologies to map the physical characteristics of the stimuli to subjective perception. While substantial progress has been made in various sensory domains, including visual and tactile [2,3], the thermoceptive and nociceptive domains remain comparatively unexplored. Among thermo-nociceptive perceptual phenomena, the thermal grill illusion (TGI) is still not well understood. The TGI manifests as a burning sensation elicited by interlaced innocuous cold and warm stimuli on the skin [4,5]. Counterintuitively, this spatial alternation does not result in the perception of each individual stimulus independently, but instead gives rise to a new, qualitatively distinct sensation characterized by unique attributes of combined hot, cold and burning sensations [6]. The perception of the TGI is thus, set apart from the sensations elicited by the individual stimuli generating it, indicating the existence of underlying sensory mechanisms that govern thermo-nociceptive interactions [7,8].

Progress in our understanding of the TGI has been constrained by a narrow focus on predetermined stimulation parameters, as for example temperatures of 20°C and 40°C [4,5,9–16] or values that deviate by only one or two degrees Celsius from this standard set of cold and warm temperatures [17–22]. The reliance on fixed temperature combinations overlooks inter-individual variability in thermal perception and has led to an overly simplistic categorization of individuals as responders or non-responders [14,23–27], without an adequate characterization of variability in temperature parameters at which the illusion manifests. Existing studies utilized fixed sets of temperatures found a responsiveness rate as low as 20-25% [25].

Previous efforts to calibrate TGI temperatures on an individual basis involved setting these temperatures at a uniform distance from the thermal pain thresholds [24,28–33], or by asking participants to report when a gradual decrease in cold temperatures and increase in warm temperatures became painful [34]. These methods indicated large inter-individual variation in responsiveness, and estimated the response rate up to 70-80% [24,29,35], significantly enhancing the proportion of the population in which TGI can be studied. The substantial differences in the response rate between studies using fixed vs. individually tailored temperatures indicates the importance of accounting for unique thermal sensitivity profiles when assessing TGI.

While previous TGI calibration methods offer some advantage over the fixed approaches, they still suffer from some limitations. The assumption of a uniform relationship between the thresholds for pain and for TGI may not be valid for all individuals. Further, the use of dynamic temperature changes when assessing pain thresholds does not accurately mimic the temporal characteristics of TGI, where stimulation temperatures are maintained constant for an extended period, from several seconds up to minutes.

To address the notable absence of adaptive psychophysical investigations of TGI perception, we developed and validated an adaptive thresholding method. This method delineates a participant-specific Thermal Grill Psychometric Function (TGPF) within two dimensions defined by cold and warm temperatures. This psychophysical calibration of the TGPF offers several advantages: it establishes a standardized and automated procedure, requiring only 30 to 40 minutes for completion, and enables the assessment of individual differences in TGI thresholds without relying on dichotomous classifications such as responders vs. non-responders. Most significantly, this approach allows more precise, rapid, and efficient estimates of this perceptual illusion in terms of probability of occurrence, perceived intensity and quality of the sensation elicited from freezing cold to burning hot.

By introducing this adaptive thresholding procedure, we characterized the psychophysical basis of the TGI and provided a method for estimating individual variability in the magnitude of the illusion. The 2D-TGC offers unique opportunities to evaluate thermo-nociceptive function in both basic research in healthy participants, and for investigating patients with altered thermal sensitivity.

## Results

We developed an adaptive thresholding procedure aimed at accurately identifying combinations of cold and warm temperatures that elicit the distinct burning sensation characteristic of the Thermal Grill Illusion (TGI). A total of 43 healthy participants completed the procedure. In each trial, we presented a stimulus, composed of three warm and two cold interlaced zones (Fig 1). Following each stimulation, participants were prompted to provide a binary response regarding their perception of a burning sensation. Subsequently, they were asked to classify their sensation as predominantly warm or cold (Fig 1). Across 100 trials, the warm and cold stimulation temperatures were adaptively adjusted to converge on specific probability levels that elicited a burning sensation. From these data, we derived the TGPF, detailing the relationship between specific warm and cold temperature combinations and the corresponding probabilities of reporting a burning sensation (Fig 2).

**Fig 1.**
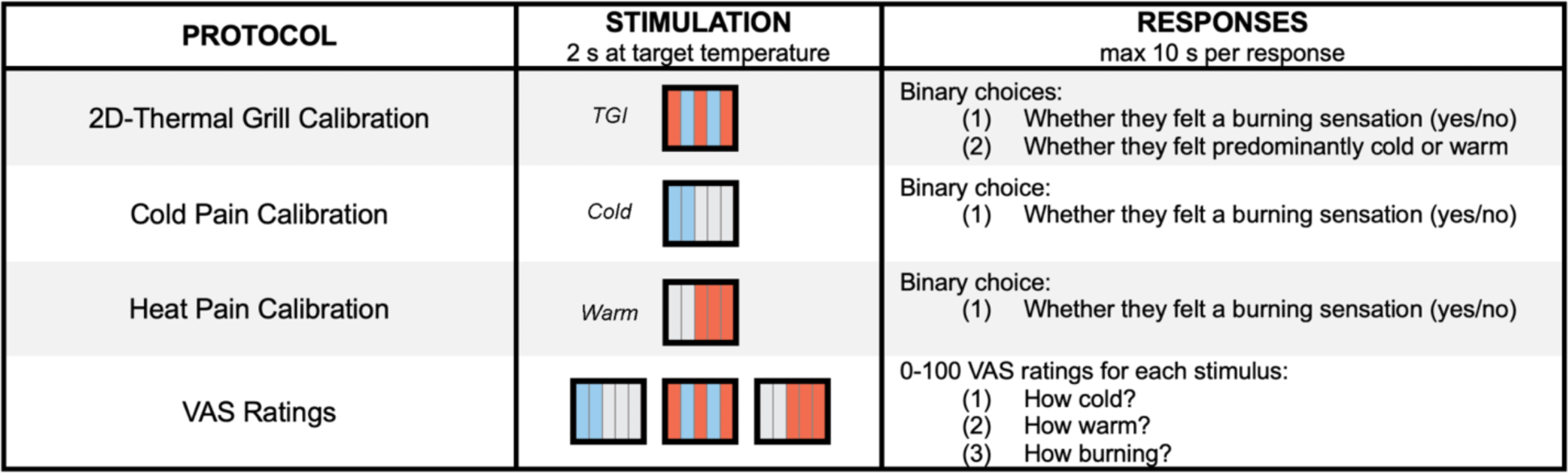
Experimental protocols and response parameters in the assessment of TGI perception. In the 2D-Thermal Grill Calibration participants received stimuli composed of interlaced warm (depicted in red) and cold (depicted in blue) zones for 2 seconds at the target temperature. They provided binary responses indicating the presence or absence of a burning sensation, and their perception as predominantly warm or cold. In the Cold Pain Calibration, stimuli consisting solely of cold zones were presented, and participants responded with a binary choice regarding the presence of a burning sensation. In the Heat Pain Calibration, stimuli consisting solely of warm zones were presented, and participants similarly responded with a binary choice about the sensation of burning. For validation purposes, participants provided VAS ratings for specific combinations of cold and warm temperature eliciting TGI and non-TGI control stimuli, which consisted of either a single cold or a single warm temperature. Following each stimulus presentation, participants rated the intensity of their cold, warm, and burning sensations using three independent VAs scales, presented sequentially.

**Fig 2.**
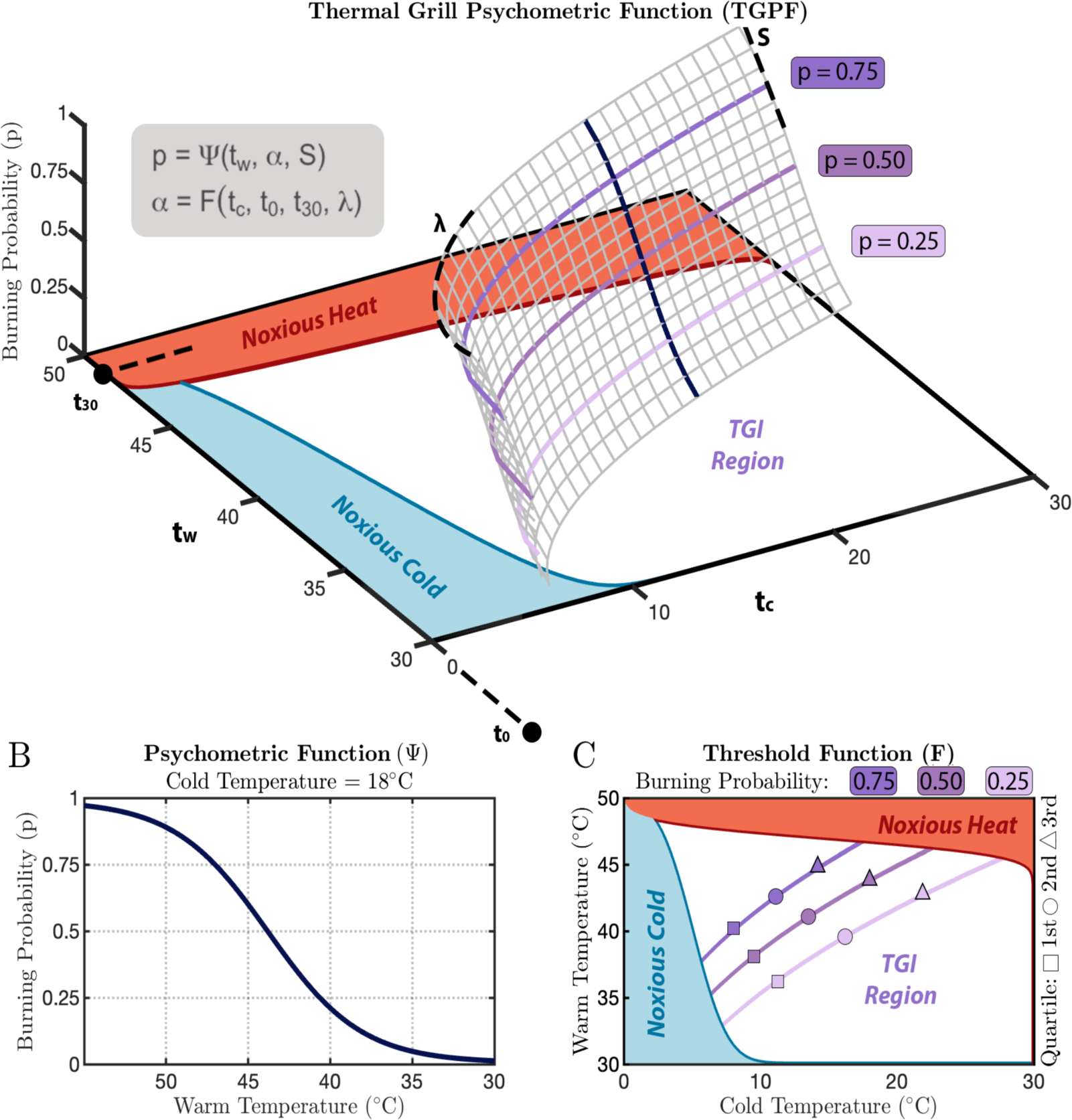
Illustration of the two-dimensional thermal grill calibration (2D-TGC) method. **(A)** The Thermal Grill Psychometric Function (TGPF) is characterized by two independent functions. **(B)** The first function (Ψ), depicted in black, represents an invariant psychometric function, mapping the probability of reporting a burning sensation from 0 to 1 at different cold and warm temperatures. **(C)** The second function (F), illustrated in purple, delineated how the perception of burning changes depending on different cold and warm temperatures. Through this approach, we successfully identified several pairs of cold and warm temperatures that were isoprobable, meaning that they induced a burning sensation with an equivalent likelihood at 25%, 50% and 75% probability levels. The boundaries of these isoprobable curves were defined by unimodal cold, depicted by the blue area labeled as “Noxious Cold”, and heat pain thresholds, depicted by the red area labeled as “Noxious Heat”. We thus effectively mapped a 3D surface, indicated as “TGI region”. Temperature combinations selected for VAS ratings are represented using three separate symbols, indicating the 1st (squares), 2nd (circles) and 3rd (triangles) quartiles of the TGI range at three specified probability levels of reporting a burning sensation. This protocol provides a comprehensive framework for assessing the TGI, which can be instrumental in both research and clinical settings.

In addition to TGI thresholding, we developed an adaptive thresholding procedure using the Psi method to assess unimodal cold and heat pain psychometric functions. The trial structure and timing were made as similar as possible to the TGI thresholding procedure. Participants received stimuli, composed of either two cold or three warm adjacent zones, and were asked to decide whether the sensation elicited was burning (Fig 1). Throughout the trials, the stimulation temperature was dynamically adjusted based on the participants’ previous responses, thus optimizing the efficiency and accuracy of threshold estimation.

To validate our TGI thresholding approach, we selected three temperature pairs for each of the estimated probability levels of 25%, 50% and 75% chance of eliciting a burning sensation, within the range of temperature defined at the two extremes by unimodal pain psychometric functions. For each stimulus, participants rated the intensity of their perceived cold, warm and burning sensations using three sequential Visual Analog Scales (VAS) (Fig 1). This step ensured that our method reliably identified temperatures that, when combined, produce the characteristic perceptual effects of the TGI, a phenomenon not observed when the individual warm and cold temperatures are presented separately.

### Characterization of the Thermal Grill Psychometric Function (TGPF) using an adaptive psychophysical method

We identified combinations of cold and warm temperatures that evoked a burning sensation by estimating the TGPF, using an adaptation of an established two-dimensional method [36]. Formally, the TGPFs were defined by two independent functions (Equation 1): a logistic psychometric function (Ψ) with steepness S, which maps pairs of temperatures to the probability of a burning sensation (p); and a threshold function F, which characterizes how the perception of burning varies with different combinations of cold (tc) and warm (tw) temperatures. The latter had three additional parameters, t0, t30, and λ, totalling four free parameters. Specifically, t0 corresponds to tw when tc is equal to zero (0°C), t30 to tw when tc is equal to the baseline temperature (30°C), and λ captures the curvature of the threshold function. These parameters were interactively estimated using a Bayesian approach.

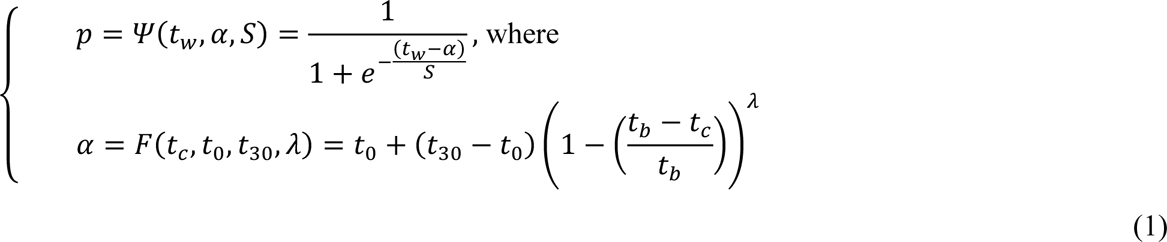

### The Thermal Grill Illusion is amenable to psychophysical thresholding

To demonstrate the feasibility of quantitatively assessing the TGI using psychophysical thresholding, we examined inter-individual variations of the estimated parameters after 100 trials (Fig 3A). For each parameter we calculated the population mean ± standard error of the mean, by bootstrapping the individual level means and standard errors. The distribution of the parameters displayed a varied range of values, reflecting individual variability in TGI sensitivity: t0 (32.1±11.3), t30 (49.4±7), λ (1.5±1.5) and S (4.1±2.7). Further, we investigated how trial-by-trial parameter estimates and their uncertainties evolved over trials (Fig S1-S2) and observed that the parameters’ standard error consistently declined over the time course of the protocol, signifying increased precision in parameter estimates.

**Fig 3.**
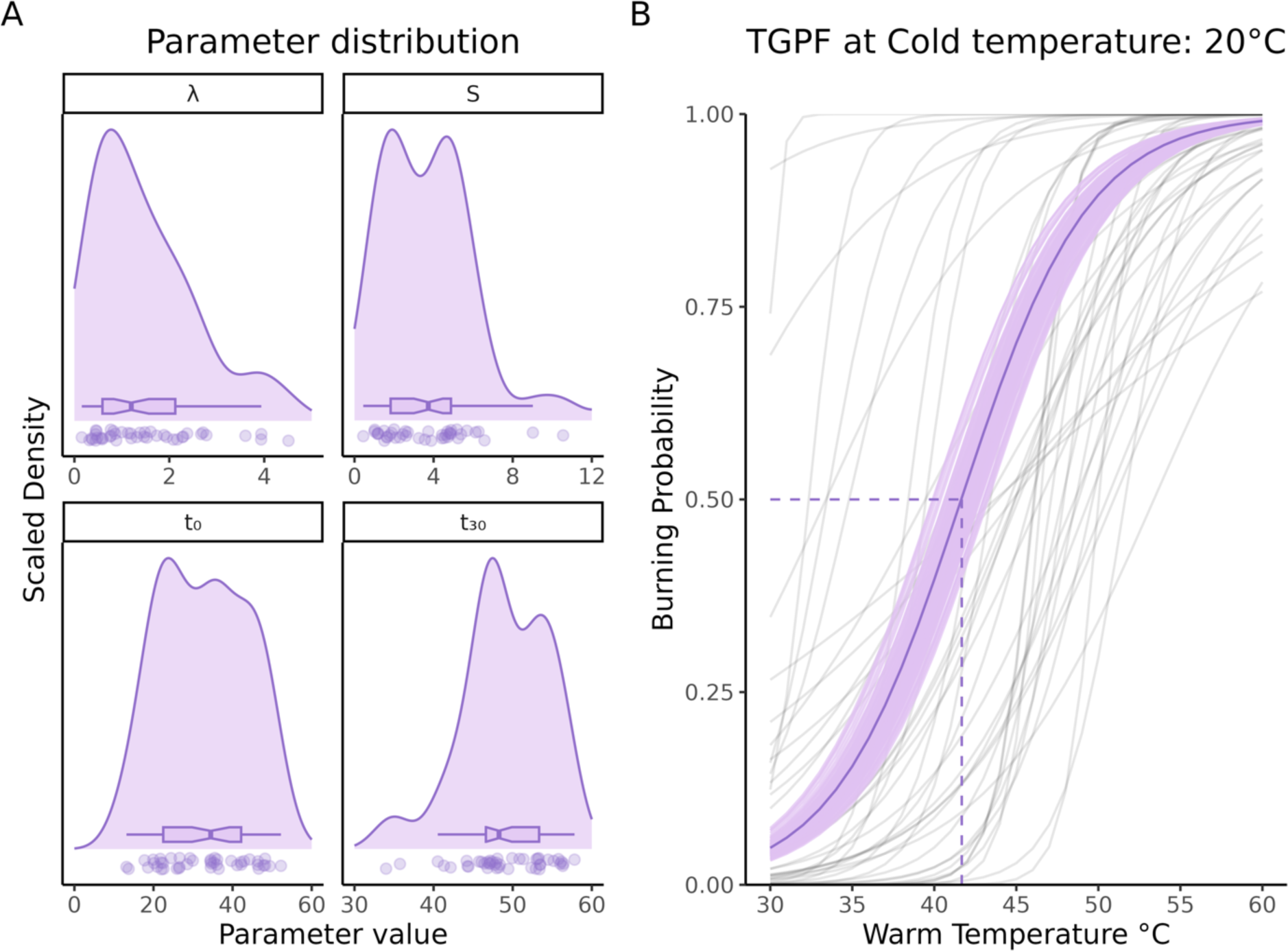
Detailed analysis of the Thermal Grill Psychometric Function (TGPF). **(A)** Population distribution of the four parameters defining the TGPF. Dots represent individual data points. **(B)** Group average and individual psychometric functions defining the relationship between a set cold temperature of 20°C and the estimated warm temperatures required to elicit TGI, in the probability range from 0 to 1. Individual participants are represented by light gray lines, while the group mean estimates are depicted by a thick purple line. The shaded purple area around the population mean represents the 95% confidence interval, obtained using bootstrapping. The dashed line represents the group mean warm temperature (41.7°C) required to elicit TGI with a 50% probability, when the cold temperature is 20°C. The 95% confidence interval around this mean corresponds to [40.6°C; 43.0°C]. Overall, these results indicate considerable variations in the cold-warm temperature pairs needed to elicit TGI sensations across individuals and challenge the use of one-size-fits-all temperature stimuli in TGI assessment.

To elucidate individual differences in TGI sensitivity, we displayed the group average alongside individual psychometric functions that estimated the probability of burning sensations given a cold stimulus of 20°C (Fig 3B). The group average indicated a 50% probability of burning sensations at the intersection of 20°C and 41.7°C, with a 95% confidence interval obtained through bootstrapping between 40.6 and 43°C. Individual psychometric functions showed large inter-individual differences in TGI sensitivity, with some participants reporting burning sensations at lower warm temperatures, while others required higher temperatures for similar sensations. This challenges the use of one-size-fits-all temperature stimuli in TGI assessments and underscores the need for individualized calibration.

In Fig 4, we presented the TGPF fits for N = 16 representative participants. While the three colored curves represent temperature combinations eliciting a burning sensation with 25%, 50% and 75% probability, the dashed black curve indicated the estimated boundary for reporting sensations as predominantly cold or warm. Individual participants’ stimulation parameters were represented by dots. These results corroborated the notion of considerable inter-individual variability in TGI sensitivity, not only in terms of probability of occurrence, but also concerning the thermosensory qualia associated with the illusion.

**Fig 4:**
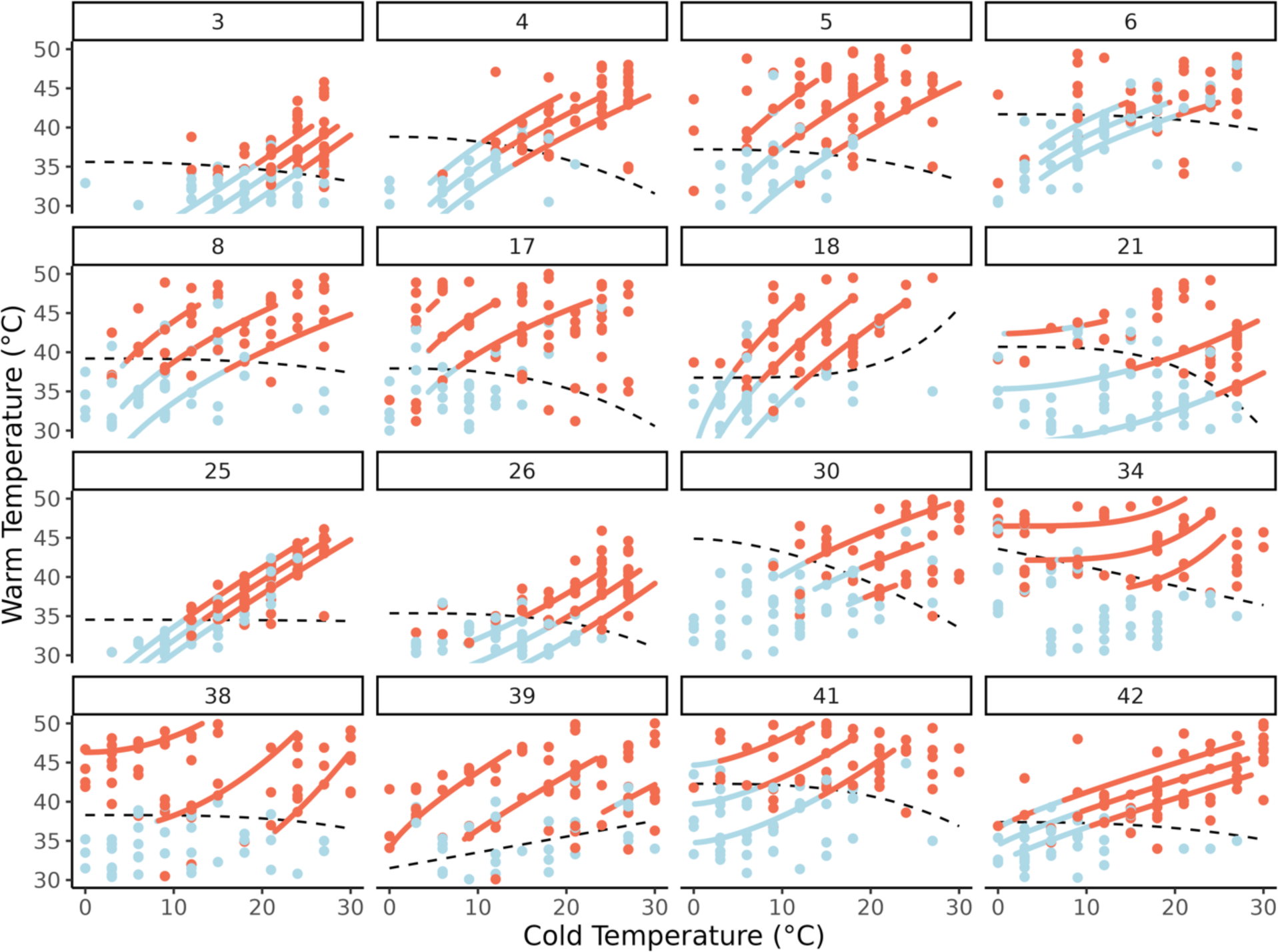
Thermal Grill Psychometric Functions (TGPF) for a subset of representative participants (N = 16). Each panel corresponds to an individual participant, with dots representing temperature pairs administered during the 2D-TGC procedure. The color of each dot reflects the binary response (cold/warm) provided by the participant. Blue dots indicate that participants selected cold, while red indicates that participants selected warm as a descriptor for their perceived thermosensory quality. The three colored curves indicate the estimated curves that elicited burning sensations with 25%, 50% and 75% probability (from bottom to top). The curves were color-coded to reflect the probability with which participants categorized the elicited sensations at a given temperature pair as either cold (blue) or warm (red). This was performed by weighting each point based on its euclidean distance from the curve. Dashed black lines represent the demarcation between cold vs. warm TGI sensations, after refitting the model using as input the quality of the TGI percept, instead of the presence/absence of burning. The two methods (euclidean distance and model refitting) provided highly similar outputs. Overall, these results indicate that it is feasible to identify several cold and warm temperature combinations that elicit TGI, with different levels of probability, and perceived thermosensory quality.

### Definition of the boundaries of the Thermal Grill Illusion

The TGPF enabled us to define a comprehensive three-dimensional surface, mapping temperature combinations that induce burning sensations at varying probability levels. However, these mappings extended across both innocuous and noxious ranges of temperatures. In the context of the TGI, the perception of burning arises from the spatial alternation of warm and cold temperatures that, when presented individually, are typically perceived as innocuous, in line with their objective physical qualities.

To precisely delineate the temperature range that effectively evokes TGI, we conducted separate assessments to determine the psychometric functions for cold and heat pain (Fig 1). The protocol followed a design similar to that of the 2D-TGC but using a one-dimensional Psi approach with stimuli that were either cold or warm. Through this thresholding, we identified specific cold and warm temperatures that were perceived as burning in isolation. To select the temperature combinations that evoked TGI, we used cold pain psychometric functions to define the lower boundary and heat pain psychometric functions to define the upper boundary, effectively demarcating a TGI region within the three-dimensional surface (Fig 2).

The parameters characterizing the pain psychometric functions were threshold (group mean ± standard error of the mean: α = 5.3±6.1 for cold and α = 45.2±2.7 for heat) and slope (β = 0.9±1.9 for cold and β = 0.7±0.9 for heat; Figure S3-4).

### Probability of occurrence, magnitude, and quality of TGI perception

To ensure the validity of our procedure, participants received nine distinct temperature combinations determined by their TGPF, and were asked to complete VAS ratings, focusing on their perceptions of cold, warm, and burning sensations. This approach was designed to verify whether the identified temperatures consistently induced the intended TGI in each individual. The nine temperature combinations of interest were equally divided across three probabilities levels, which corresponded to the 25%, 50% and 75% likelihood of reporting a burning sensation. Within each of these probability levels, we selected three temperature combinations, which corresponded to the first, second and third quartiles within the range demarcated by cold and heat pain psychometric functions at the two extremes. By selecting these three quartiles, we aimed to capture a comprehensive range of temperature values that could be used to evoke the illusion (Figure 2). In addition to using cold and warm temperature combinations for assessing the TGI, we presented a set of unimodal thermal stimuli. This included nine exclusively cold and nine exclusively warm temperatures, at values determined by the TGPF. However, unlike the combined stimuli used for TGI assessment, these temperatures were presented in isolation. These stimuli provided a baseline for comparison against the participants’ responses to the combined stimuli and were necessary to evaluate the participants’ responsivity to the illusion.

VAS ratings were analyzed using mixed effects zero one inflated beta regressions. Analyses on the beta distribution (i.e., continuous values greater than 0 and less than 1) revealed that combinations of temperatures (i.e., TGI stimuli) were rated as more intensely burning than any of the corresponding unimodal stimuli (Figure 5). Specifically, sensations of burning were overall significantly more intense during TGI, compared to unimodal warm stimuli (*β* = -0.38, 95% CI = [-0.57; -0.18], p < .0001), or unimodal cold stimuli (*β* = -0.26, 95% CI = [-0.44; -0.07], p < .05). For all types of stimuli, the burning intensity increased with increasing burning probability levels (*β* = 1.66, 95% CI = [1.44; 1.88], p < .0001) and this increase was more pronounced (steeper slope) for TGI than unimodal cold (*β* = -0.78, 95% CI = [-1.12; -0.44], p < .0001) or warm stimuli (*β* = -0.47, 95% CI = [-0.83; -0.11], p < .05).

**Fig 5:**
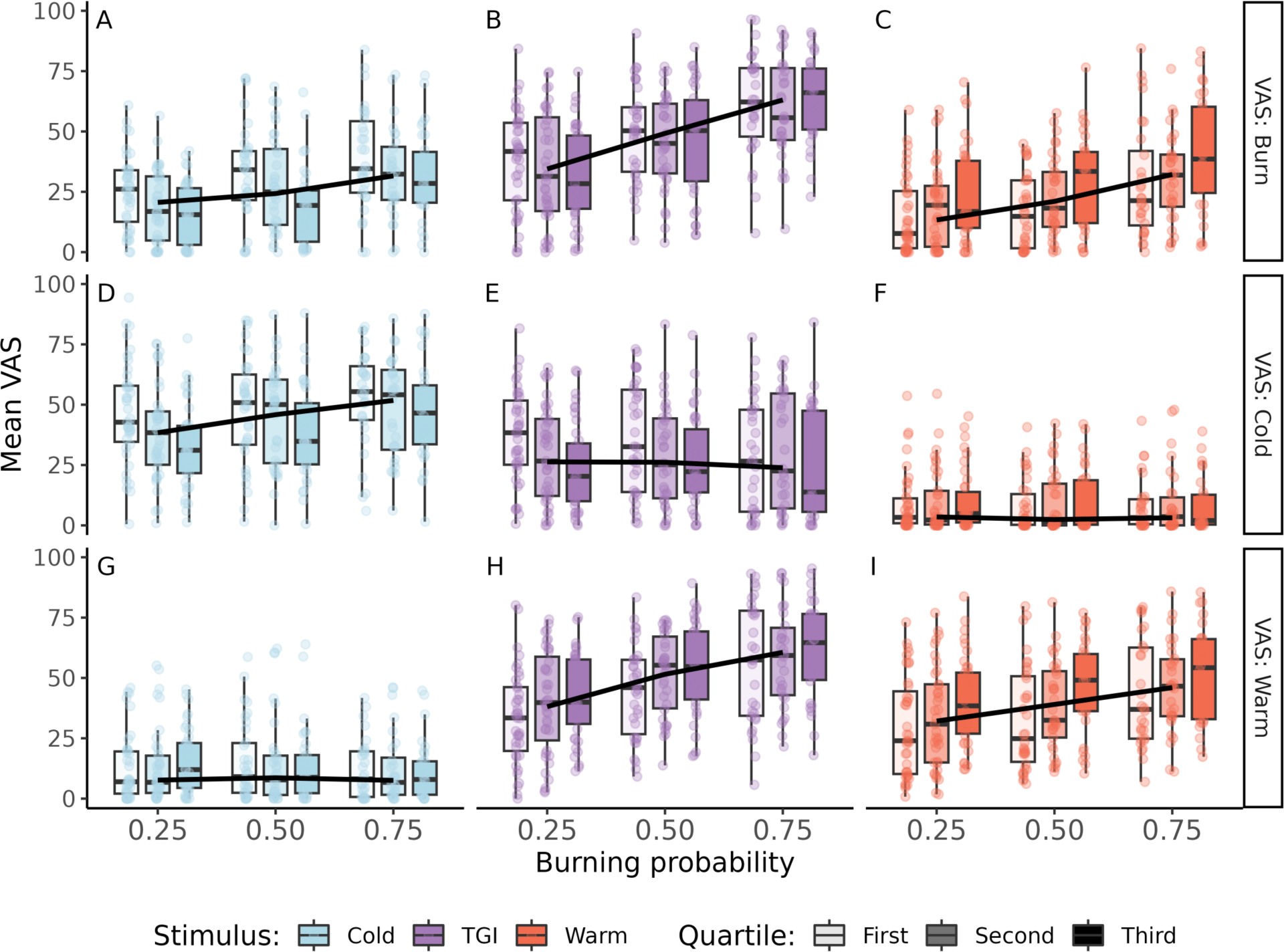
Overview of subjective ratings for TGI and non-TGI (cold and warm) control stimuli. Mean VAS ratings (burn, cold, and warm) across different stimulus types: cold (A, D, G), TGI (B, E, H) and warm (C, F, I), and across three burning probabilities: p = 0.25, p = 0.50 and p = 0.75. Individual participant means are indicated by small dots and black lines connect the median value of the three burning probability levels. We show that the TGI was perceived as more burning than individual cold (A vs. B) and warm (B vs. C) stimuli. Further, the TGI was perceived as less cold than individual cold (D vs. E), but warmer than individual warm (H vs. I) stimuli. Data points from isoprobable curves, corresponding to the three probability levels of reporting a burning sensation, were indicated by three variations of the same color. Results showed that burning and warm ratings increased with increasing burning probability. However, distinct temperature pairs, belonging to the same isoprobable curve, elicited burning and warm sensations of similar intensity.

Sensations of warmth were similar for TGI compared to unimodal warm (*β* = -0.08, 95% CI = [-0.23; 0.08], p = 0.33). Warm ratings also significantly increased as a function of burning probability levels for TGI (*β* = 1.70, 95% CI = [1.49; 1.91], p < .0001) and, to a lower extent for unimodal warm (*β* = -0.60, 95% CI = [-0.89; -0.30], p < .0001). Further, sensations of cold were similar during TGI and unimodal cold (*β* = 0.10, 95% CI = [-0.06; 0.27], p = 0.21). Cold ratings significantly increased with increasing burning probability for cold stimuli (*β* = 0.78, 95% CI = [0.46; 1.09], p < .0001), but not for TGI stimuli (*β* = -0.03, 95% CI = [-0.28; 0.22], p = 0.81).

These findings are in line with the conceptualization of the TGI as reflecting heat enhancement, accompanied by burning or painful sensations. This clear pattern of responses validated the effective elicitation of the TGI using TGPF, and demonstrated how the selection of distinct temperatures effectively shapes the illusion in accordance with varying probability levels. In sum, we demonstrated the feasibility of TGI calibration by applying principles of psychophysics.

### Assessment of the thermosensory qualia of TGI perception

While we revealed a modulation of the intensity of burning sensations across three distinct burning probability levels, the TGI burning sensation was remarkably consistent within each probability level (Fig 5). For temperature combinations within the same isoprobable curve, corresponding to the first, second, and third quartiles within the range demarcated by cold and heat pain psychometric functions, participants reported similar burning sensations *β* = -0.03, 95% CI = [- 0.17; 0.11], p = 0.69. This finding aligns with the principles of the TGPF, indicating that a diverse range of temperature combinations can induce similar burning sensations. Despite these consistent levels of burning sensations, the perceived thermosensory quality of the TGI varied depending on the quartile from which the temperature combination was sampled. To better understand this variation in thermal perception, we computed a thermosensory index using the formula: cold rating / (cold rating + warm rating) for each stimulus type. Here, a value approaching one signifies a perception as predominantly cold, while a value closer to zero suggests a perception as predominantly warm. This ratio provides information on how individuals perceive the thermal aspect of the TGI, beyond the intensity of burning sensations alone (Fig 6).

**Fig 6.**
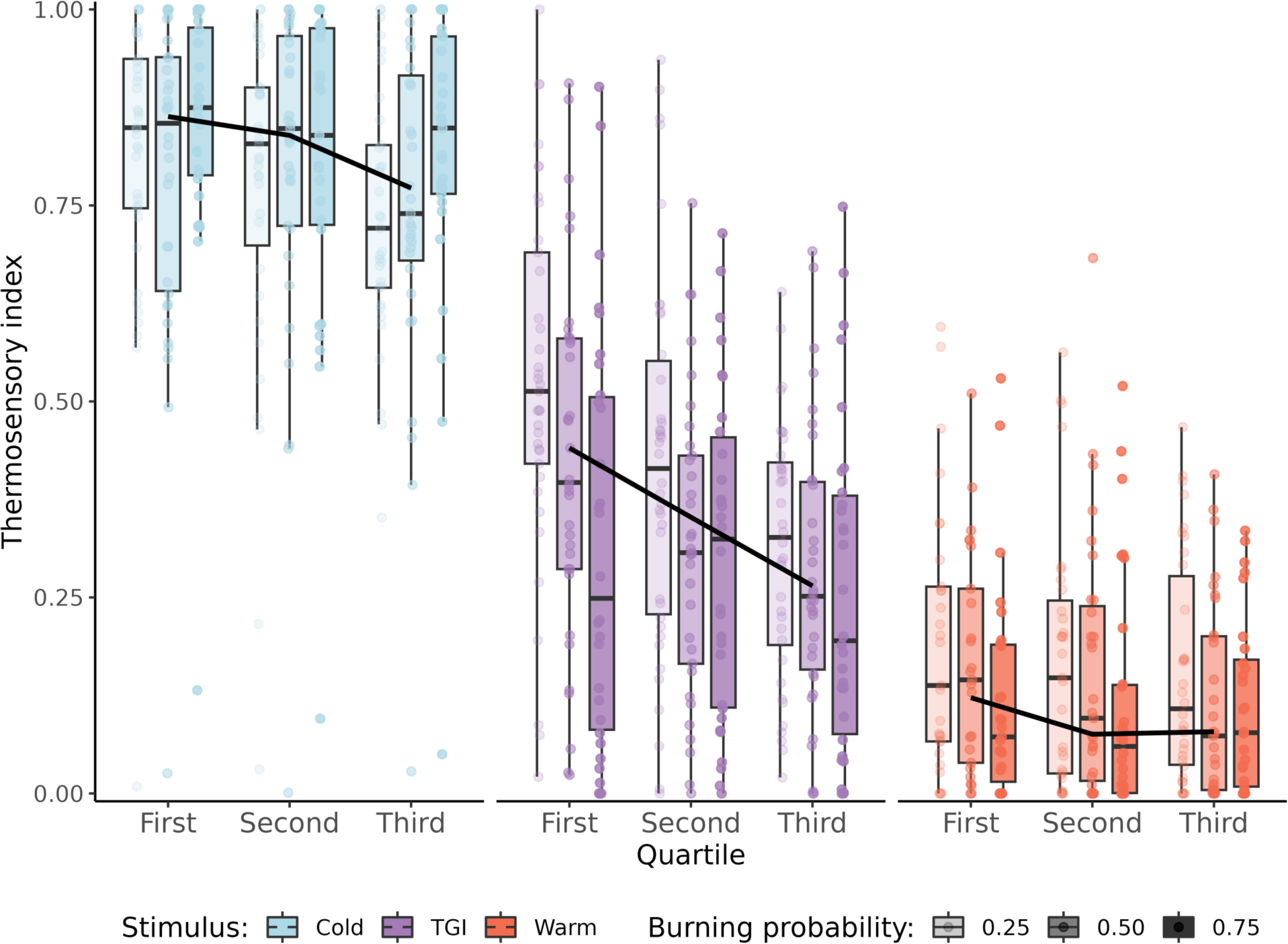
Perceived thermosensory qualities of TGI at equivalent intensity of burning sensation. TGI perception is typically characterized by burning sensations coupled with enhanced cold or warm thermosensory qualities. To quantify these thermosensory qualities in TGI perception, we derived a thermosensory index where values closer to one indicate a stronger sensation of cold, and values closer to zero indicate a more pronounced sensation of warmth. Each participant’s average perception is represented by a small dot. The results revealed significant variability in thermosensory quality among individuals for stimuli that elicited comparable levels of burning sensations. While some individuals reported TGI as primarily cold, others experienced it as predominantly warm. However, on average, the data shows a group tendency towards the perception of warmth during TGI. This warmth perception varied with the intensity of the temperature pairs, with those in the first quartile being reported as colder compared to the more intensely warm sensations reported in the second and third quartiles.

We analyzed how the thermosensory index changed based on stimulus type and quartile and found that the perceived predominant thermosensory quality was significantly more influenced by the quartile for TGI stimuli compared to both unimodal warm *β* = 0.48, 95% CI = [0.08; 0.88], p < .05, and unimodal cold stimuli *β* = 0.47, 95% CI = [0.10; 0.84], p < .05. This result indicated that as we moved along the same equiprobable curve, we observed larger variations in the perceived thermosensory quality of the TGI stimuli, compared to when the unimodal stimulus was either cold or warm. In addition, a main effect of burning probability indicated that with increasing burning probability, the perceived thermosensory quality became predominantly warmer, irrespective of stimulus type *β* = -0.46, 95% CI = [-0.62; -0.30], p < .0001.Overall, these results indicate that it is in principle possible to identify temperature combinations that elicit TGI with distinct thermosensory attributes, from cold to warm, within the same individual.

### Classification of TGI responsivity along a continuum

TGI responsivity is defined by the positive difference between burning ratings for temperature pairs vs. individual temperature stimuli. However, previous research has largely defined TGI responsivity as a dichotomous attribute (i.e., responders vs. non-responders) based on how participants responded to a unique set of temperature, most commonly 20 and 40°C. The TGPF allows for the characterization of TGI responsivity as a continuous index.

To analyze TGI responsivity, we rescaled burning ratings from 0 to 1, and computed the mean burning rating for each of the three types of stimuli at each of the three burning probabilities. We then subtracted the maximum burning sensation elicited by the unimodal stimuli from the mean burning sensation elicited by the TGI stimulus. This yielded a value for each participant and burning probability, ranging from -1 to 1; where negative values indicated enhanced burning sensation for the unimodal stimulus, while positive values indicated enhanced burning sensation for temperature combinations, using identical objective temperatures. A value of 0 suggested no difference between the burning ratings when stimuli were unimodal or temperature pairs. Using uncertainty propagation, we derived a 95% confidence interval for each participant and burning probability for the responsivity index. The resulting distribution in our sample is depicted in Fig 7, sorted from lowest to highest responsivity index within each burning probability. When considering the responsivity index for temperatures eliciting TGI with 50% and 75% probability, only one participant had a confidence interval overlapping zero, indicating a lack of responsivity, while two participants had an on opposite pattern of responsivity (i.e., greater burning sensation for the unimodal temperature vs. combined cold and warm temperatures).

**Fig 7.**
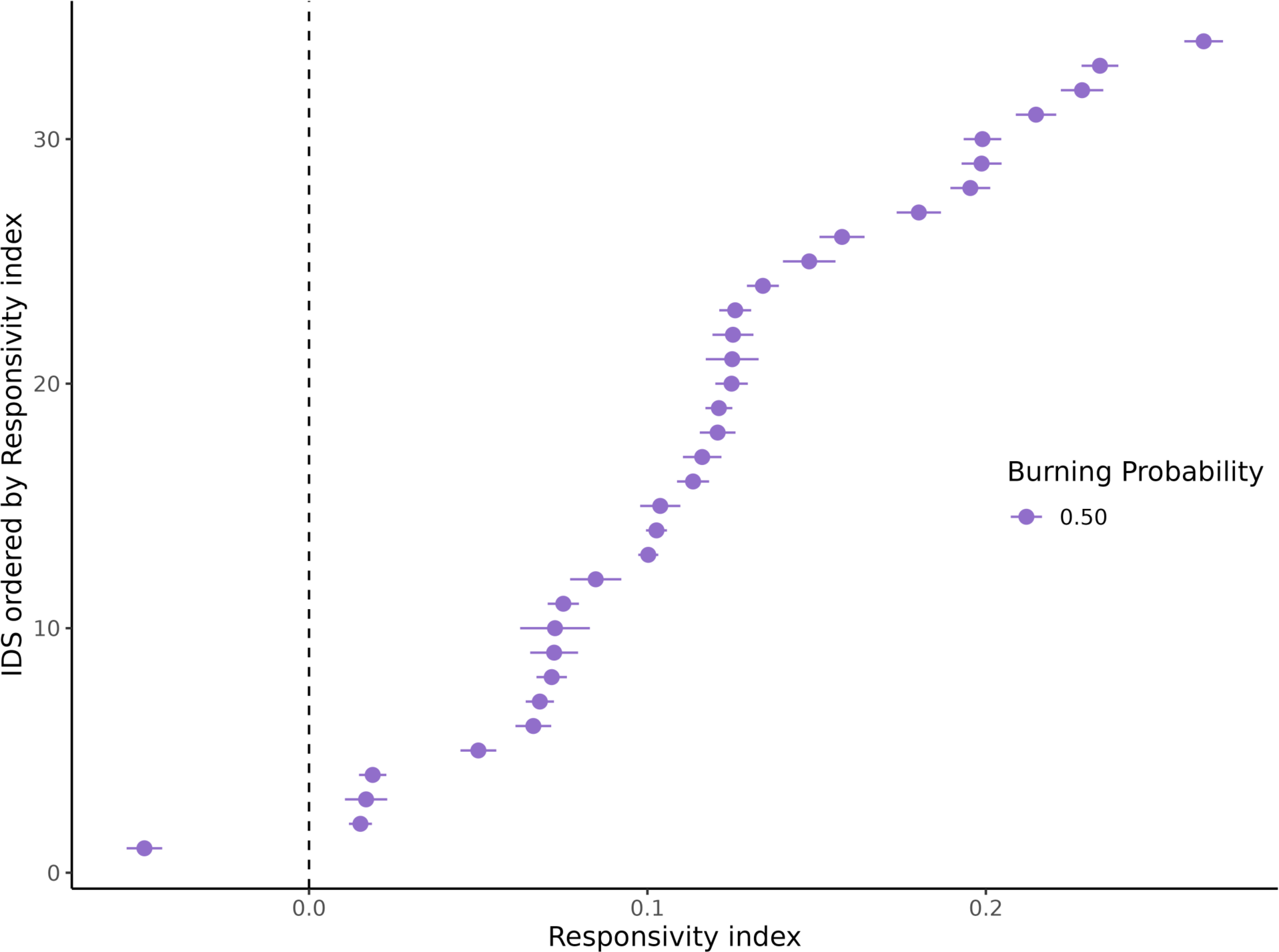
Responsivity index conditional on 50% burning probability. The figure displays the responsivity index for the burning probability of p = 0.50. The responsivity index is calculated as the difference between the mean burning rating of either the unimodal cold or warm (the highest is selected) and the burning rating elicited by the temperature combinations. Positive values indicate higher burning perception during TGI compared to the unimodal stimulus, 0 means no difference in the burning sensation across stimulus typed, and negative values indicate higher burning sensations for the unimodal stimulus compared to the temperature pairs. Using uncertainty propagation, we calculated the 95% confidence interval for the responsivity index of each participant. The y-axis represents participant IDs; however, the order of the participant IDs is rearranged based on the magnitude of the thermosensory index (from low to high). This analysis demonstrates how TGI responsivity can be characterized as a continuous variable.

## Discussion

This study has established the Thermal Grill Illusion (TGI) as a quantifiable phenomenon through the application of psychophysical methods. By introducing the Two-Dimensional Thermal Grill Calibration (2D-TGC) protocol, we provided a psychophysical framework capable of delineating a range of temperature combinations that effectively induce TGI. This method can estimate varying probabilities of experiencing the illusion, and pinpoint distinct temperature combinations eliciting varying degrees of perceived intensity and thermosensory quality, thus accounting for the considerable variability in how this phenomenon is perceived by different individuals.

Validation of the 2D-TGC protocol was achieved using Visual Analog Scale (VAS) ratings from participants, confirming the successful elicitation of the illusion through the spatial alternation of cold and warm temperatures, as opposed to presenting each temperature individually. Participants reported burning sensations of similar magnitude across all temperature combinations along the same isoprobable curve, corresponding to the probabilities of 25%, 50%, and 75% for reporting burning sensations. This demonstrated that a variety of temperature combinations can equally provoke the illusion. Further, we showed large inter-individual variability with respect to the required warm temperature when the cold temperature was set at 20°C, indicating the inadequacy of choosing fixed temperatures, such as 20°C and 40°C, to assess the TGI [4,5,9–16]. In particular, using our specific thermode, 20°C needed to be paired with a temperature between 40.6 and 43°C to elicit the illusion with 50% probability in most individuals.

One unique advantage of this approach is its capability of quantitatively characterizing two perceptual dimensions inherent to the TGI, related to illusory pain and temperature enhancement. Characteristically, the TGI manifests as pronounced burning sensations, accompanied by enhanced perception of warmth or cold [6–8,35]. We showed that burning sensations increased in magnitude depending on the specific isoprobability curve (increasing from 25% to 50% and 75%) and that thermosensory qualities varied depending on both the quartile (increased heat progressing from the first to the second and the third quartiles) and the improbability curve (increased heat progressing from 25% to 50% and 75%) from which the temperature pairs were sampled. These results indicate that the TGI can manifest with a broad spectrum of intensity and thermosensory qualities within the same individual. Our approach thus quantifies not only between-subject variability in this phenomenon, but also captures within-subject variations in how the illusion is experienced across a comprehensive range of values.

Previous research aiming to calibrate TGI temperatures based on individual pain thresholds found an association between the magnitude of the TGI and the difference between cold and warm stimuli, suggesting that larger temperature differentials correlate with a higher likelihood of experiencing the illusion [24,28]. The 2D-TGC method enhances this understanding by precisely defining the threshold function F, which quantifies the impact of temperature differentials on the probability of perceiving the illusion. Furthermore, our approach incorporated a parameter-optimization procedure that customizes these temperature differentials for each individual, thereby accommodating and fine-tuning the protocol to reflect each participant’s unique sensitivity to the burning sensation.

Overall, the 2D-TGC provides an efficient and precise tool for TGI assessment, with key implications for future experimental and clinical research. We successfully applied the TGPF across all individuals in our study, extracting meaningful parameters related to TGI perception for the entire sample under investigation. This approach supersedes the traditional dichotomous classification of participants into responders or non-responders based on static, arbitrary temperature thresholds - a practice previously commonplace in the field, (e.g., [14,20,26]). As a result, our approach addresses common challenges associated with participant recruitment by eliminating the need to exclude non-responders at the screening stage or to limit analysis to a subset of responders. Further, it allows for the assessment of TGI responsivity on a continuum, rather than a dichotomous attribute, offering a granular quantification of the TGI phenomenon across multiple perceptual features.

The ability to adaptively titrate the TGI through psychophysical staircasing is particularly relevant for clinical studies involving between-group comparisons. Previous studies investigating whether TGI is enhanced or reduced in specific clinical populations failed to account for the inter-individual differences in thermal sensitivity (e.g., [21,22]). Given the proposed role of TGI as a model for chronic pain in conditions such as central neuropathic [37,38] and nociplastic pain [11,29], adopting an approach that captures the entire spectrum of individual responses to TGI is crucial for elucidating its significance in clinical settings.

A key limitation of our modeling approach is the assumption of a fixed slope across the surface defined by the TGPF. An improved version would account for changes in slopes, especially at the edges of the TGI region. We expect that steeper slopes towards the heat pain boundary and flatter curves towards the cold pain boundary may provide a more accurate representation of the space of temperatures eliciting this illusion. However, an increased number of trials would be necessary for estimating varying slopes. Nevertheless, we provide evidence for the feasibility of thresholding procedures to capture different features of a perceptual phenomenon related to thermo-nociceptive interactions. This approach can be combined with neurophysiological or neuroimaging techniques to better understand the underlying mechanisms of TGI and can be applied in clinical studies to assess the relationship between TGI and chronic pain in diverse populations. Further, this multi-dimensional approach can be potentially adapted to investigate other thermal and pain phenomena, such as paradoxical heat sensations - a phenomenon similar to the TGI, but involving the temporal alternation of warming and cooling on the same skin area [39–41].

In summary, we provided a principled psychophysical approach to calibrate the probability of TGI occurrence, and to quantify TGI responses across a comprehensive spectrum of perceptual states. This spectrum includes the entire range of sensations, from the intensity of burning sensations to the perception of the illusion on a continuum between freezing cold and burning heat. Our findings highlight the importance of adopting a multi-dimensional psychophysical framework for TGI assessment, facilitating an accurate representation of inter-individual differences in temperature and pain perception. Our method can be used in future studies to investigate illusions of pain in various clinical conditions, particularly those associated with chronic pain.

## Materials and method

### Participants

The study involved 43 healthy individuals, across two experiments. The first experiment included 28 participants (age = 25.3 ± 5.6, 14 males), while the second experiment included 15 participants (age = 26.3 ± 6.6, 5 males). Across both experiments we maintained consistent inclusion and exclusion criteria. Participants were physically and psychologically healthy, as determined by the absence of any major neurological, psychiatric or pain-related disorder, and could not have made use of any analgesic medications or recreational drugs, either regularly or in the week leading up to the experiment. Prior to the inclusion in the study, all participants received comprehensive briefings regarding the research procedures and provided their informed consent. The study adhered to the ethical guidelines outlined in the Declaration of Helsinki and received approval from the local Institutional Review Board at the Danish Neuroscience Center at Aarhus University.

### Stimuli

All thermal stimuli were delivered using a Thermal Cutaneous Stimulator (TCS II, QST.Lab), comprising five distinct stimulation zones. Stimuli designed to elicit the TGI consisted of three zones set to warm temperatures and two zones to cold temperatures, arranged in an alternated spatial pattern. Unimodal stimuli consisted of either two adjacent zones set to cold temperatures, or three adjacent zones set to warm temperatures only. The remaining zones were maintained at a baseline temperature of 30°C (Fig 1). The two experiments involved the use of two distinct TCS II probes. In experiment 1, each stimulation zone measured 7 x 28mm, covering a surface area of 10 cm2, and the nominal rate of temperature change of the probe was 20°C/s. In experiment 2, each stimulation zone measured 7.4 x 24.2 mm, resulting in a total surface area of 9 cm2, and the nominal rate of temperature change of the probe was 100°C/s. In both experiments, each stimulus was maintained at target temperature for 2 seconds, before the temperature of all five zones returned to baseline (30°C). The inter-trial interval, before the next stimulus presentation, lasted between 2 and 4 seconds. The thermode was repositioned every 3 trials to one of three possible locations on the volar forearm, to minimize potential habituation or sensitization effects. The computerized task was created with the Psychophysics Toolbox Version 3 (PTB-3) [42].

### Adaptive staircasing: Estimating TGI thresholds

For staircasing of the TGI, we developed a two-dimensional adaptive procedure to identify combinations of cold and warm temperatures that elicited burning sensations. This staircase was implemented using the Functional Adaptive Sequential Testing (FAST) toolbox [36] (https://github.com/edvul/FAST-matlab) in MATLAB (R2020a). TThe FAST toolbox enables the adaptive testing of multiple dimensions simultaneously by augmenting the mathematical characterization of the psychometric function (Ψ), relating a given performance level on a task not only to the intensity of a single independent variable and the steepness of the curve, but also by considering the influence of a mediating variable. The relationship between the independent variable and the mediating variable is mathematically described by the threshold function F. This function can take many shapes and is characterized by an array of parameters, which are estimated experimentally.

This approach is constrained by three assumptions: i) the psychometric function (Ψ) translates independently across the threshold function F; ii) the slope is constant for points along the threshold function F; and iii) other parameters of the psychometric function (lapse and guess rates) are fixed. The algorithm is initiated with priors for each parameter and employs Bayes’ theorem to update the posterior probability distribution iteratively, ensuring that the prior for each trial is the previous trial’s posterior. The FAST toolbox can be flexibly applied in various sensory systems and experimental conditions, requiring three main definitions tailored to each specific setup: the configuration of the threshold function F’s equation, a sampling strategy, and a criterion for optimization.

In this study, we modeled the probability of experiencing burning sensations (p) using a logistic psychometric function (Ψ) with inverse steepness S, as a function of cold (tc) and warm (tw) temperatures. These cold and warm temperatures were linked by the threshold function F as detailed in Equation 1. The priors for t0 and t30 were specified as Gaussian distributions, N(30,10) and N(50,10), respectively. Meanwhile, the parameters λ and S were assigned uniform distributions within the ranges 0 - 4 and 0.001 - 20, respectively. The adaptive procedure determined which temperatures to present for each trial using the following sampling strategy. First, the algorithm randomly selected a cold temperature (tc) within the range between 0°C to 27°C (Experiment 1) and 0°C to 30°C (Experiment 2), with discrete intervals of 3°C. Second, it dynamically adjusted the warm temperatures (tw) to aim for a specific response probability (25%, 50%, or 75% levels), assigned randomly for each trial. To do so, it determined the most likely values for each free parameter (posterior modes) and used them to determine the warm temperature needed to achieve the target burning probability. This optimization process was repeated, starting from the random selection of a cold temperature, if the selected tw exceeded 50°C or was below the baseline temperature (tb).

### Adaptive staircasing: Estimating thermal pain thresholds

For adaptive staircasing of cold and heat pain thresholds, we used the Psi method, implemented via the Palamedes toolbox [43] in MATLAB (R2020a). Psi is a Bayesian adaptive algorithm designed for efficient estimation of psychometric function parameters, specifically the threshold (location parameter) and slope (rate-of-change parameter) [44]. It operates by iteratively updating a posterior probability distribution on these parameters based on each response of the participant, to optimize stimulus placement for threshold and slope estimations. A similar approach was previously used to estimate the threshold and slope of psychometric functions for cold and warm detection [45,46].

In Experiment 1, individual pain thresholds were estimated based on 30 stimuli, using a logistic psychometric function and uniform priors. The heat pain threshold prior was set within the range of 35°C to 50°C, while the cold pain one spanned from 0°C to 29°C. The slope prior ranged from 0.001 to 6. The guess rate was set to 0, and the lapse rate was set to 0.1. In Experiment 2, each pain threshold was estimated based on 50 stimuli, using a cumulative normal psychometric function. The heat pain threshold prior was uniform in the 35°C to 50°C range, while for cold pain it spanned from 0°C to 30°C. The value of the slope prior ranged from 0.001 to 6. The guess and lapse rates were both set to 0.05.

Additionally, in Experiment 2 we introduced 10 demonstration trials before cold and warm pain threshold assessments, with the aim to familiarize the participants with the procedure and with different stimulation intensities. The intensities of these trials were determined using an up-down staircase, with step size decreasing as a function of the number of reversals. For cold pain, the initial temperature value was set at 20°C, and the initial downward step size was 5°C. The downward step size gradually decreased at each reversal at a rate of 1°C, and was coupled with a consistent upward step size of 2°C. Cold temperature cut-offs were 0°C and 30°C. For heat pain, the initial temperature value was set at 38°C, the initial upward step size was 3°C, diminishing at a rate of 1°C at each reversal, while the downward step size was fixed at 2°C. Warm temperature cut-offs were 35°C and 50°C.

### Refining the boundaries of the three dimensional TGI surface

While we defined a three-dimensional surface describing cold-warm temperature pairs that were associated with specific probabilities of burning perception for each individual (i.e., the TGPF), these temperatures were not limited to the innocuous thermosensory range. We thus constrained the boundaries of the surface post hoc, based on the pain thresholds obtained using the Psi procedure, to define a TGI region where combinations of innocuous temperatures trigger an illusion of pain. Importantly, the temperatures presented to the participants were constrained within a range between minimum 0oC for cold temperatures and maximum 50oC for warm. In Fig 4, the lower and upper bounds of each burning probability curve was determined by cold and heat psychometric functions. Thus, temperature pairs for which tc < Ψcold(p) or tw > Ψheat(p) are not displayed.

### Subjective validation of the TGPF

To validate whether we adequately induced an illusion of pain, we asked participants to provide subjective ratings reflecting their perception of TGI and non-TGI control stimuli. TGI stimuli consisted of cold and warm temperature pairs along three curves, corresponding to probabilities of 25%, 50% and 75% of reporting a burning sensation. For each of these isoprobable curves, we selected three equidistant points within the TGI range (temperature pairs at the first, second and third quartile of each curve). Non-TGI control stimuli were the same cold and warm temperatures that comprised the TGI stimuli, but presented individually and paired with the baseline temperature of 30°C. Each unique TGI and non-TGI temperature combination was repeated five times, for a total of 135 stimulation trials, presented in a randomized order. After each stimulus, participants rated the intensity of their cold, warm, and burning sensations utilizing three separate Visual Analog Scales (VAS), presented in a randomized order. The lower limit (i.e., 0) corresponded to a complete absence of a sensation, while the upper extreme (i.e., 100) indicated the most intense sensation.

To investigate the perceived thermosensory quality of TGI and non-TGI stimuli, we calculated a thermosensory index reflecting whether each stimulus was perceived as predominantly cold or warm, based on cold and warm VAS ratings. This index was computed as the ratio of cold ratings to the sum of cold and warm ratings. In cases where the variable was undefined (i.e., both cold and warm ratings were zero), the ratio was set to 0.5.

### Statistics

We analyzed perceptual variables of interest (VAS ratings, thermosensory index) using a mixed effects zero one inflated beta (ZOIB) regressions, with the logit link function, within the gamlss package in R [47]. Each VAS rating was analyzed using a separate ZOIB model (Table S1-S4). This modeling approach was appropriate to accurately account for the bounded nature of the VAS rating scale as well as the presence of zeros and ones in our data. Zero values were provided by a participant if they did not perceive any burning sensation or if the thermosensory rating contradicted the stimulation quality (e.g., cold ratings for warm stimuli and warm ratings for cold stimuli). The proportions of zeros were 24.4 ± 3.0% (mean ± standard error of the mean) for cold ratings, 24.4, 3%, 15.9, 2.2% for warm ratings and 21.4, 3.3% for burning ratings. Finally, the proportions of ones were 0.5, 0.4% for cold ratings, 0.3, 0.2% for warm ratings and 0.6, 0.3% for burning ratings. The models concerning the distribution of ones only included an intercept for each participant, due to the low proportion of data points.

The mixed effect models concerning beta and zero distributions included the main effects and interactions of interest specified in the main text and in the supplementary tables (Table S1-S3). All models included trial number and stimulus duration as covariates, to account for habituation and or sensitization effects as well as the total amount of stimulus received. The random effects structure of the models included participants nested within the two experiments to account for the use of two slightly different thermodes across Experiment 1 and 2. Between experiment variance was negligible, indicating that the results from the two experiments were highly overlapping.

## Data visualization

Data were visualized using custom scripts in matlab and R, using the ggplot2 package [48] and raincloud plots [49].

## Code and data availability statement

All code and data will be available on GitHub and OSF following acceptance of the manuscript.

## Acknowledgements

This work was supported by the European Research Council under the grants ERC-2020-StG-948838 (FF, CSD, JEF, AGM, CEK) and ERC-2020-StG-948788 (MGA, ASC). We also acknowledge funding from the Lundbeck Foundation under the grants R272-2017-4345 (MGA, NN). We thank Ed Vul for insightful comments on a preliminary version of the TGPF. We also thank Patrik Molnar and Sára Anna Szabó for help with data collection.

## Authors contributions

Author contributions listed alphabetically according to CRediT taxonomy: Conceptualization: CSD, FF. Data curation: CSD, JFE, FF. Formal analysis: CSD, JFE, FF. Funding acquisition: FF. Investigation: CSD. Methodology: ASC, CSD, JFE, FF, AGM, NN. Project administration: CSD, FF. Resources: MGA, ASC, CSD, JFE, FF, CEK, AGM. Software: ASC, CSD, JFE, FF, AGM. Supervision: FF. Visualization: CSD, JFE, FF. Writing – original draft: CSD, JFE, FF. Writing – review & editing: MGA, ASC, JFE, FF, AGM, NN.

## References

1. Nour MM, Nour JM. Perception, illusions and Bayesian inference. Psychopathology. 2015;48: 217–221. doi:10.1159/000437271

2. Eagleman DM. Visual illusions and neurobiology. Nat Rev Neurosci. 2001;2: 920–926. doi:10.1038/35104092

3. Longo MR, Haggard P. Weber’s illusion and body shape: Anisotropy of tactile size perception on the hand. J Exp Psychol Hum Percept Perform. 2011;37: 720–726. doi:10.1037/a0021921

4. Craig AD, Bushnell MC. The thermal grill illusion: Unmasking the burn of cold pain. Science. 1994;265: 252–255. doi:10.1126/science.8023144

5. Craig AD, Reiman EM, Evans A, Bushnell MC. Functional imaging of an illusion of pain. Nature. 1996;384: 258–260. doi:10.1038/384258a0

6. Bach P, Becker S, Kleinböhl D, Hölzl R. The thermal grill illusion and what is painful about it. Neurosci Lett. 2011;505: 31–35. doi:10.1016/j.neulet.2011.09.061

7. Fardo F, Finnerup NB, Haggard P. Organization of the Thermal Grill Illusion by Spinal Segments. Ann Neurol. 2018;84: 463–472. doi:10.1002/ana.25307

8. Fardo F, Beck B, Allen M, Finnerup NB. Beyond labeled lines: A population coding account of the thermal grill illusion. Neurosci Biobehav Rev. 2020;108: 472–479. doi:10.1016/j.neubiorev.2019.11.017

9. Averbeck B, Rucker F, Laubender RP, Carr RW. Thermal grill-evoked sensations of heat correlate with cold pain threshold and are enhanced by menthol and cinnamaldehyde. Eur J Pain. 2013;17: 724–734. doi:10.1002/j.1532-2149.2012.00239.x

10. Averbeck B, Seitz L, Kolb FP, Kutz DF. Sex differences in thermal detection and thermal pain threshold and the thermal grill illusion: A psychophysical study in young volunteers. Biol Sex Differ. 2017;8: 29. doi:10.1186/s13293-017-0147-5

11. Bäumler P, Brenske A, Winkelmann A, Irnich D, Averbeck B. Strong and aversive cold processing and pain facilitation in fibromyalgia patients relates to augmented thermal grill illusion. Sci Rep. 2023;13: 15982. doi:10.1038/s41598-023-42288-7

12. Horing B, Kerkemeyer M, Büchel C. Temporal Summation of the Thermal Grill Illusion is Comparable to That Observed Following Noxious Heat. J Pain. 2023;0. doi:10.1016/j.jpain.2023.11.015

13. Hunter J, Dranga R, Wyk M van, Dostrovsky JO. Unique influence of stimulus duration and stimulation site (glabrous vs. Hairy skin) on the thermal grill-induced percept. Eur J Pain. 2015;19: 202–215. doi:10.1002/ejp.538

14. Jutzeler CR, Warner FM, Wanek J, Curt A, Kramer JLK. Thermal grill conditioning: Effect on contact heat evoked potentials. Sci Rep. 2017;7: 40007. doi:10.1038/srep40007

15. Leung A, Shukla S, Li E, Duann J-R, Yaksh T. Supraspinal characterization of the thermal grill illusion with fMRI. Mol Pain. 2014;10: 18. doi:10.1186/1744-8069-10-18

16. Matsuda S, Igawa Y, Uchisawa H, Iki S, Osumi M. Thermal Grill Illusion in Post-Stroke Patients: Analysis of Clinical Features and Lesion Areas. J Pain Res. 2023;16: 3895–3904. doi:10.2147/JPR.S433309

17. Harper DE, Hollins M. Coolness both underlies and protects against the painfulness of the thermal grill illusion. Pain. 2014;155: 801–807. doi:10.1016/j.pain.2014.01.017

18. Harper De, Hollins M. Conditioned pain modulation dampens the thermal grill illusion. Eur J Pain. 2017;21: 1591–1601. doi:10.1002/ejp.1060

19. Li X, Petrini L, Wang L, Defrin R, Arendt-Nielsen L. The importance of stimulus parameters for the experience of the thermal grill illusion. Neurophysiol Clin. 2009;39: 275–282. doi:10.1016/j.neucli.2009.06.006

20. Lindstedt F, Lonsdorf TB, Schalling M, Kosek E, Ingvar M. Perception of Thermal Pain and the Thermal Grill Illusion Is Associated with Polymorphisms in the Serotonin Transporter Gene. PLoS One. 2011;6: e17752. doi:10.1371/journal.pone.0017752

21. Sumracki NM, Buisman-Pijlman FTA, Hutchinson MR, Gentgall M, Rolan P. Reduced response to the thermal grill illusion in chronic pain patients. Pain Med. 2014;15: 647–660. doi:10.1111/pme.12379

22. Kim HC, Chang MC, Oh SH, Lee SB, Yang SY, Shin DA. Thermal Grill Illusion in Chronic Lower Back Pain: A Case-Control Study. J Pain Res. 2023;16: 1573–1579. doi:10.2147/JPR.S403387

23. Bekrater-Bodmann R, Chung BY, Richter I, Wicking M, Foell J, Mancke F, et al. Deficits in pain perception in borderline personality disorder: Results from the thermal grill illusion. Pain. 2015;156: 2084–2092. doi:10.1097/j.pain.0000000000000275

24. Bouhassira D, Kern D, Rouaud J, Pelle-Lancien E, Morain F. Investigation of the paradoxical painful sensation (’illusion of pain’) produced by a thermal grill. Pain. 2005;114: 160–167. doi:10.1016/j.pain.2004.12.014

25. Schaldemose EL, Horjales-Araujo E, Svensson P, Finnerup NB. Altered thermal grill response and paradoxical heat sensations after topical capsaicin application. Pain. 2015;156: 1101. doi:10.1097/j.pain.0000000000000155

26. Scheuren R, Sütterlin S, Anton F. Rumination and interoceptive accuracy predict the occurrence of the thermal grill illusion of pain. BMC Psychol. 2014;2: 22. doi:10.1186/2050-7283-2-22

27. Scheuren R, Duschek S, Schulz A, Sütterlin S, Anton F. Blood pressure and the perception of illusive pain. Psychophysiology. 2016;53: 1282–1291. doi:10.1111/psyp.12658

28. Adam F, Alfonsi P, Kern D, Bouhassira D. Relationships between the paradoxical painful and nonpainful sensations induced by a thermal grill. Pain. 2014;155: 2612–2617. doi:10.1016/j.pain.2014.09.026

29. Adam F, Jouët P, Sabaté J-M, Perrot S, Franchisseur C, Attal N, et al. Thermal grill illusion of pain in patients with chronic pain: A clinical marker of central sensitization? Pain. 2023;164: 638–644. doi:10.1097/j.pain.0000000000002749

30. Kern D, Pelle-Lancien E, Luce V, Bouhassira D. Pharmacological dissection of the paradoxical pain induced by a thermal grill. Pain. 2008;135: 291–299. doi:10.1016/j.pain.2007.12.001

31. Kern D, Plantevin F, Bouhassira D. Effects of morphine on the experimental illusion of pain produced by a thermal grill. Pain. 2008;139: 653–659. doi:10.1016/j.pain.2008.07.001

32. Caston RM, Davis TS, Smith EH, Rahimpour S, Rolston JD. A novel thermoelectric device integrated with a psychophysical paradigm to study pain processing in human subjects. J Neurosci Methods. 2023;386: 109780. doi:10.1016/j.jneumeth.2022.109780

33. Schaldemose EL, Raaschou-Nielsen L, Böhme RA, Finnerup NB, Fardo F. It is one or the other: No overlap between healthy individuals perceiving thermal grill illusion or paradoxical heat sensation. Neurosci Lett. 2023;802: 137169. doi:10.1016/j.neulet.2023.137169

34. Defrin R, Benstein-Sheraizin A, Bezalel A, Mantzur O, Arendt-Nielsen L. The spatial characteristics of the painful thermal grill illusion. Pain. 2008;138: 577–586. doi:10.1016/j.pain.2008.02.012

35. Mitchell AG, Ehmsen JF, Christensen DE, Stuckert A, Haggard P, Fardo F. Disentangling the spinal mechanisms of illusory heat and burning sensations in the Thermal Grill Illusion. bioRxiv; 2023. doi:10.1101/2023.08.24.554485

36. Vul E, Bergsma J, MacLeod DIA. Functional adaptive sequential testing. Seeing Perceiving. 2010;23: 483–515. doi:10.1163/187847510x532694

37. Craig BAD. Can the basis for central neuropathic pain be identified by using a thermal grill? Pain. 2008;135: 215–216. doi:10.1016/j.pain.2008.01.022

38. Rosner J, Andrade DC de, Davis KD, Gustin SM, Kramer JLK, Seal RP, et al. Central neuropathic pain. Nat Rev Dis Primers. 2023;9: 1–19. doi:10.1038/s41572-023-00484-9

39. Mitchell AG, Ehmsen JF, Basińska M, Courtin AS, Böhme RA, Deolindo CS, et al. Temporal Contrast Enhancement in Thermosensation: A Framework for Understanding Paradoxical Heat Sensation. bioRxiv; 2023. doi:10.1101/2023.08.23.554435

40. Schaldemose EL, Andersen NT, Finnerup NB, Fardo F. When cooling of the skin is perceived as warmth: Enhanced paradoxical heat sensation by pre-cooling of the skin in healthy individuals. Temperature. 2023;10: 248–263. doi:10.1080/23328940.2022.2088028

41. Vollert J, Fardo F, Attal N, Baron R, Bouhassira D, Enax-Krumova EK, et al. Paradoxical heat sensation as a manifestation of thermal hypesthesia: A study of 1090 patients with lesions of the somatosensory system. Pain. 2024;165: 216–224. doi:10.1097/j.pain.0000000000003014

42. Kleiner M, Brainard D, Pelli D, Ingling A, Murray R, Broussard C. What’s new in psychtoolbox-3. Perception. 2007;36: 1–16.

43. Prins N, Kingdom FAA. Applying the Model-Comparison Approach to Test Specific Research Hypotheses in Psychophysical Research Using the Palamedes Toolbox. Front Psychol. 2018;9: 1250. doi:10.3389/fpsyg.2018.01250

44. Kingdom FAA, Prins N. Psychometric Functions∗. In: Kingdom FAA, Prins N, editors. Psychophysics (Second Edition). San Diego: Academic Press; 2016. pp. 55–117. doi:10.1016/B978-0-12-407156-8.00004-9

45. Courtin AS, Slootjes SM, Caty G, Hermans MP, Plaghki L, Mouraux A. Assessing thermal sensitivity using transient heat and cold stimuli combined with a Bayesian adaptive method in a clinical setting: A proof of concept study. Eur J Pain. 2020;24: 1812–1821. doi:10.1002/ejp.1628

46. Courtin AS, Delvaux A, Dufour A, Mouraux A. Spatial summation of cold and warm detection: Evidence for increased precision when brisk stimuli are delivered over larger area. Neurosci Lett. 2023;797: 137050. doi:10.1016/j.neulet.2023.137050

47. Rigby RA, Stasinopoulos DM. Generalized additive models for location, scale and shape. J R Stat Soc, C: Appl Stat. 2005;54: 507–554. doi:10.1111/j.1467-9876.2005.00510.x

48. ggplot2: Elegant Graphics for Data Analysis (3e). Available: https://ggplot2-book.org/

49. Allen M, Poggiali D, Whitaker K, Marshall TR, Langen J van, Kievit RA. Raincloud plots: A multi-platform tool for robust data visualization. Wellcome Open Res. 2019;4: 63. doi:10.12688/wellcomeopenres.15191.2

