## Supplementary Material for "Assessing Individual Sensitivity to the Thermal Grill Illusion: A Two-Dimensional Adaptive Psychophysical Approach"

### Supplementary Figures

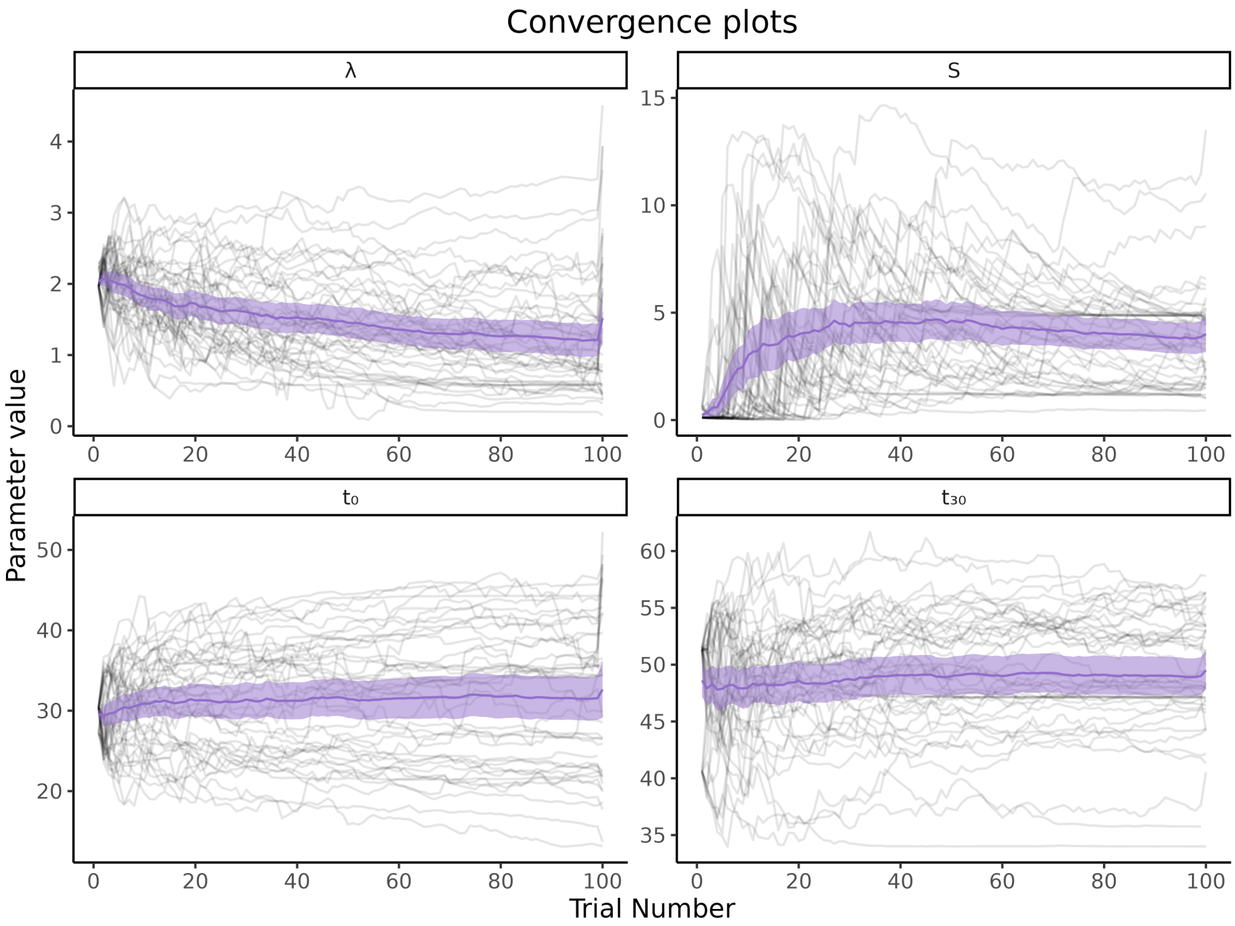


**Fig S1. Trial-by-trial estimates for each parameter characterizing the TGPF.** Each participant is represented by a light gray line, while the thick violet line depicts the group mean estimate on each trial, and the shaded purple lines the 95% confidence interval.

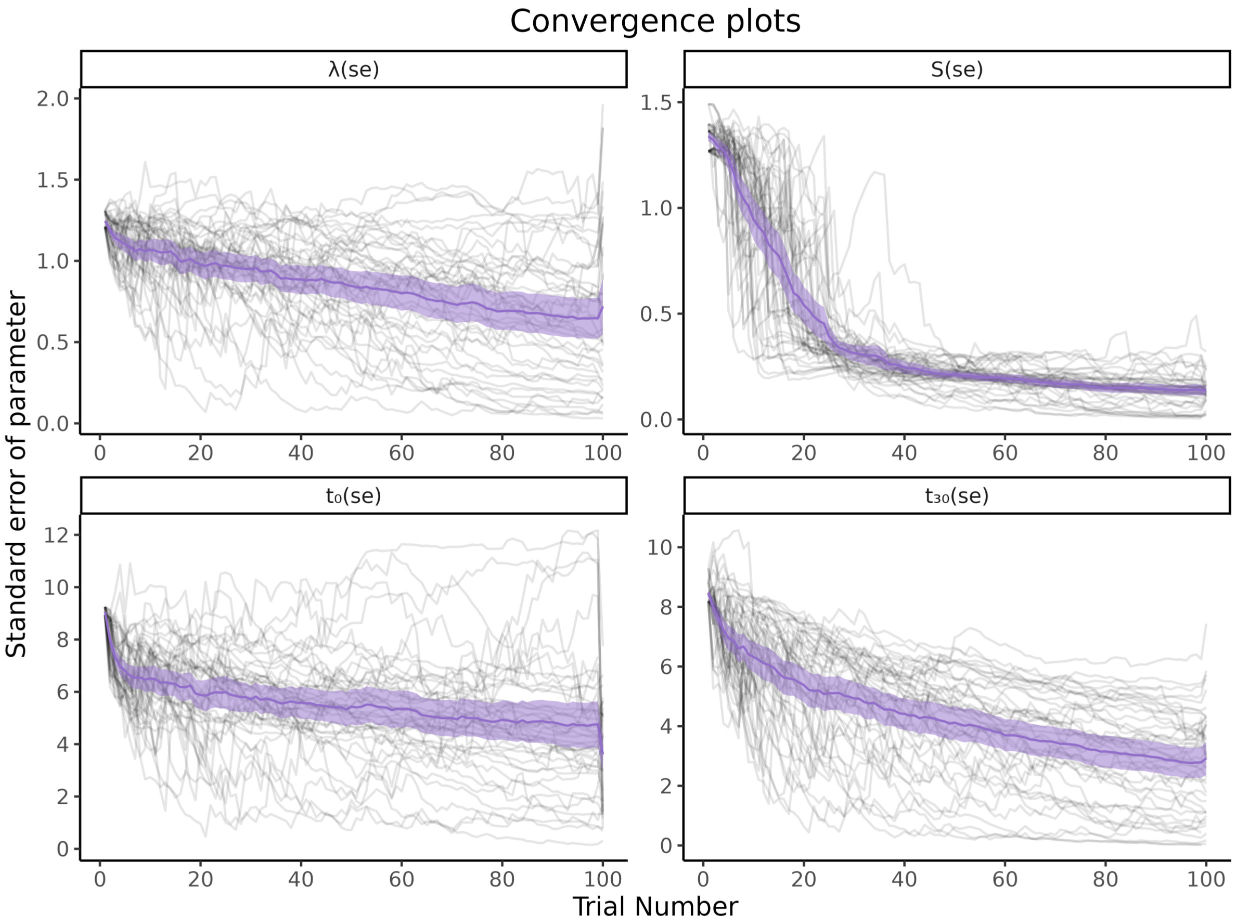


**Fig S2. Staircase convergence of TGPFs parameters in our sample of N = 43 individuals.** Staircase convergence of all parameters, computed as the posterior dispersion for all participants (mean and 95% confidence interval in purple), indicating a discernible and steep decrease over the course of the experiment. Each participant is represented by a light gray line.

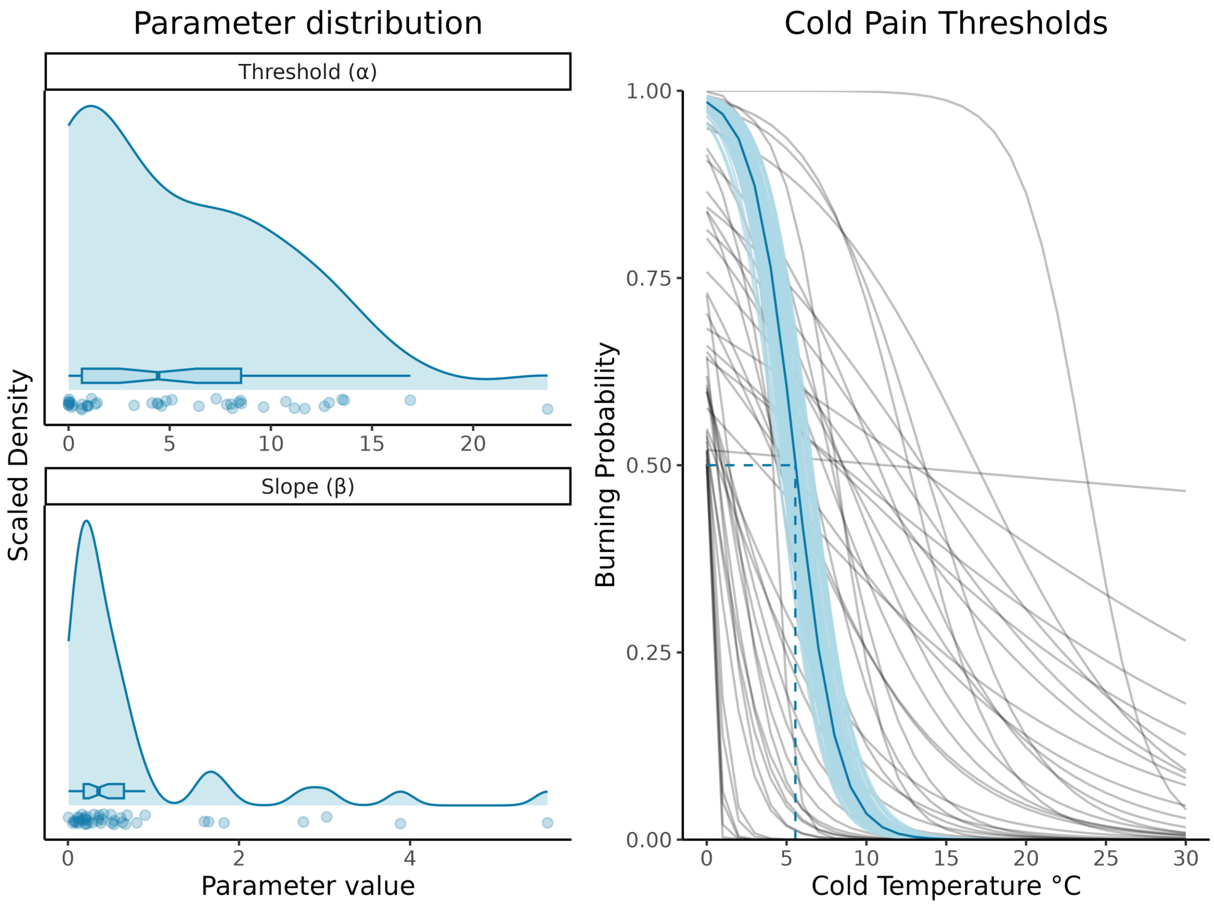


**Fig S3. Cold Pain Psychometric Functions.** Distribution of the threshold and slope parameters, as well as pain psychometric functions for each individual and at the group-level. The shaded area around the group-level PF indicates the 95% confidence interval. The dashed line represents the group mean cold temperature (5.6°C) required to elicit pain with a 50% probability. The 95% confidence interval around this mean corresponds to [4.8°C ; 6.2°C] obtained through bootstrapping.

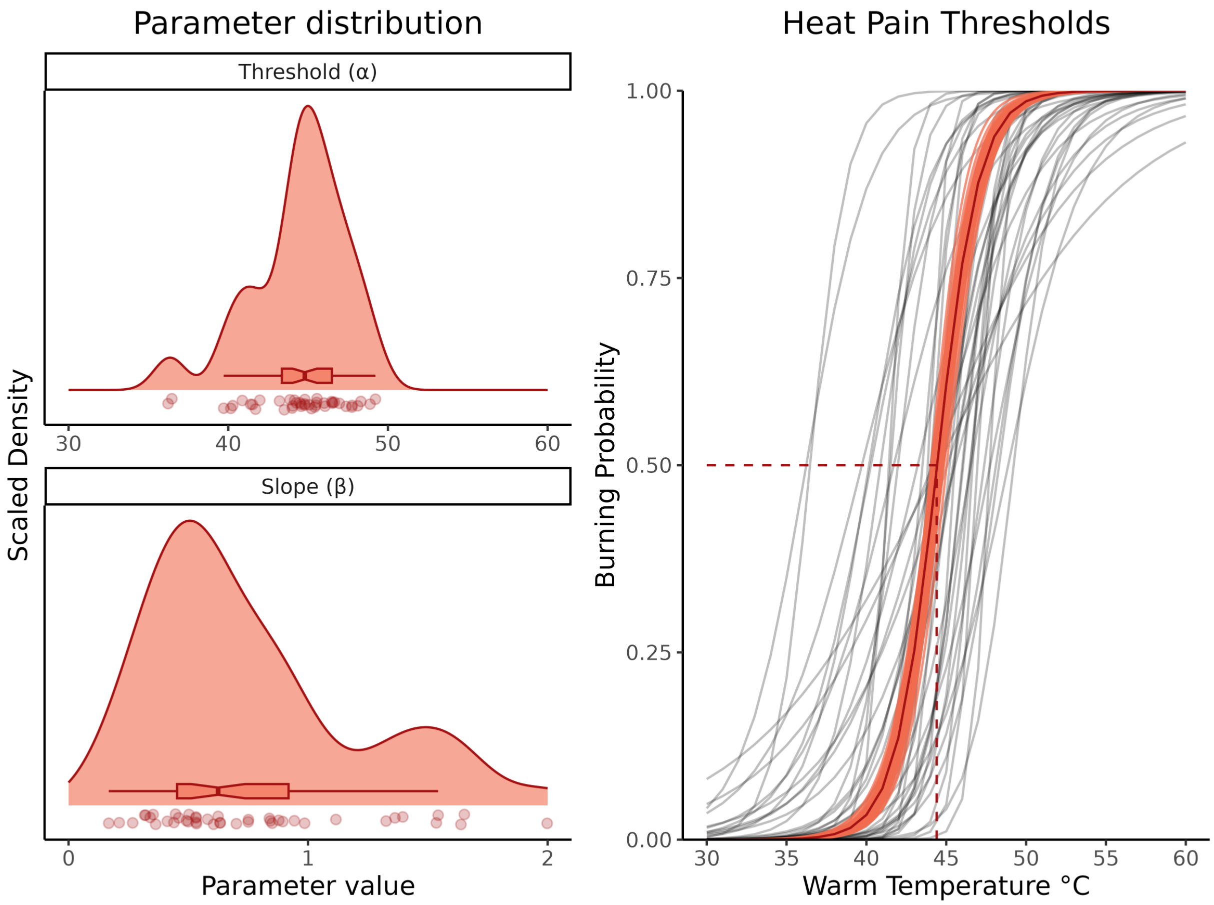


**Fig S4. Heat Pain Psychometric Functions.** Distribution of the threshold and slope parameters, as well as pain psychometric functions for each individual and at the group-level. The shaded area around the group-level PF indicates the 95% confidence interval. The dashed line represents the group mean warm temperature (44.4°C) required to elicit pain with a 50% probability. The 95% confidence interval around this mean corresponds to [44.1°C ; 44.8°C] obtained through bootstrapping.

#### Supplementary Tables

Supplementary Tables (S1-S4) present the formulation of four distinct models: cold, warm, burning VAS ratings, and the thermosensory index. For each model, fixed and random effects are specified in the first row. We report all parameters (, , and ) of each mixture model (i.e., the zero-one inflated beta regression), as well as the contrasts for all fixed effects. The parameters are as follows: is the mean of the beta distribution, is standard deviation of the beta distribution, is the proportion of zero from a Bernoulli distribution and lastly is the proportion of ones, also from a Bernoulli distribution.

*Cold Ratings*

| VAS_Cold~Stimulus * Burning_prob + Quartile + Trial_n + Stim_dur + re(random = ~1 \| experiment/ID), ZOIB(link = logit) | | | | | |
|---|---|---|---|---|---|
| **parameter** | **contrast** | **β** | **SE** | **t** | **p** |
| μ | (Intercept) | -0.5 | 0.091 | -5.5 | 4.4e-08 |
| μ | StimulusCold | 0.1 | 0.083 | 1.2 | 0.21 |
| μ | StimulusWarm | -0.88 | 0.11 | -7.7 | 1.8e-14 |
| μ | Burning_prob | -0.03 | 0.13 | -0.24 | 0.81 |
| μ | Quartile | -0.54 | 0.072 | -7.5 | 8.2e-14 |
| μ | Trial_n | -0.0021 | 0.00038 | -5.5 | 4.3e-08 |
| μ | Stim_dur | 0.071 | 0.011 | 6.5 | 9.2e-11 |
| μ | StimulusCold:Burning_prob | 0.78 | 0.16 | 4.9 | 1.2e-06 |
| μ | StimulusWarm:Burning_prob | -0.14 | 0.22 | -0.63 | 0.53 |
| σ | (Intercept) | -0.37 | 0.093 | -4 | 7.2e-05 |
| σ | StimulusCold | -0.084 | 0.091 | -0.92 | 0.36 |
| σ | StimulusWarm | -0.13 | 0.11 | -1.2 | 0.24 |
| σ | Burning_prob | -0.078 | 0.13 | -0.58 | 0.56 |
| σ | Quartile | -0.042 | 0.077 | -0.54 | 0.59 |
| σ | Trial_n | -9.9e-05 | 0.0004 | -0.25 | 0.8 |
| σ | Stim_dur | 0.032 | 0.011 | 2.8 | 0.0045 |
| σ | StimulusCold:Burning_prob | -0.15 | 0.18 | -0.87 | 0.39 |
| σ | StimulusWarm:Burning_prob | -0.015 | 0.22 | -0.069 | 0.95 |
| ν | (Intercept) | -0.33 | 0.068 | -4.9 | 9.2e-07 |
| τ | (Intercept) | -5.8 | 0.27 | -22 | 6.1e-98 |

Table 1, Cold ratings

*Warm Ratings*

| VAS_Warm~Stimulus * Burning_prob + Quartile + Trial_n + Stim_dur + re(random = ~1 \| experiment/ID), ZOIB(link = logit) | | | | | |
|---|---|---|---|---|---|
| **parameter** | **contrast** | **β** | **SE** | **t** | **p** |
| μ | (Intercept) | -1.2 | 0.087 | -14 | 4.7e-43 |
| μ | StimulusCold | -0.63 | 0.1 | -6.3 | 4.5e-10 |
| μ | StimulusWarm | -0.077 | 0.078 | -0.98 | 0.33 |
| μ | Burning_prob | 1.7 | 0.11 | 16 | 3.8e-53 |
| μ | Quartile | 0.6 | 0.067 | 9 | 5e-19 |
| μ | Trial_n | -0.00064 | 0.00035 | -1.8 | 0.068 |
| μ | Stim_dur | 0.041 | 0.012 | 3.5 | 0.00055 |
| μ | StimulusCold:Burning_prob | -1.7 | 0.19 | -8.6 | 1.6e-17 |
| μ | StimulusWarm:Burning_prob | -0.6 | 0.15 | -4 | 6.1e-05 |
| σ | (Intercept) | -0.33 | 0.09 | -3.7 | 0.00021 |
| σ | StimulusCold | 0.093 | 0.1 | 0.9 | 0.37 |
| σ | StimulusWarm | -0.11 | 0.086 | -1.3 | 0.18 |
| σ | Burning_prob | 0.07 | 0.12 | 0.6 | 0.55 |
| σ | Quartile | -0.23 | 0.072 | -3.2 | 0.0014 |
| σ | Trial_n | -0.00053 | 0.00038 | -1.4 | 0.16 |
| σ | Stim_dur | 0.022 | 0.012 | 1.8 | 0.07 |
| σ | StimulusCold:Burning_prob | -0.28 | 0.2 | -1.4 | 0.17 |
| σ | StimulusWarm:Burning_prob | 0.041 | 0.16 | 0.25 | 0.8 |
| v | (Intercept) | -0.69 | 0.54 | -1.3 | 0.2 |
| v | StimulusCold | 2.9 | 0.52 | 5.5 | 3.4e-08 |
| v | StimulusWarm | 0.19 | 0.54 | 0.35 | 0.73 |
| v | Burning_prob | -4 | 0.77 | -5.3 | 1.4e-07 |
| v | Quartile | -2.8 | 0.45 | -6.2 | 7.3e-10 |

| VAS_Warm~Stimulus * Burning_prob + Quartile + Trial_n + Stim_dur + re(random = ~1 \| experiment/ID), ZOIB(link = logit) | | | | | |
|---|---|---|---|---|---|
| **parameter** | **contrast** | **β** | **SE** | **t** | **p** |
| ν | Trial_n | -0.0049 | 0.0024 | -2.1 | 0.04 |
| ν | Stim_dur | 0.0032 | 0.071 | 0.045 | 0.96 |
| ν | StimulusCold:Burning_prob | 4.6 | 1.1 | 4.3 | 1.8e-05 |
| ν | StimulusWarm:Burning_prob | -1.3 | 1.2 | -1.1 | 0.29 |
| τ | (Intercept) | -6 | 0.28 | -22 | 4.6e-98 |

Table 2, Warm ratings

*Burning Ratings*

| VAS_Burn~Stimulus * Burning_prob + Quartile + Trial_n + Stim_dur + re(random = ~1 \| experiment/ID), ZOIB(link = logit) | | | | | |
|---|---|---|---|---|---|
| **parameter** | **contrast** | **β** | **SE** | **t** | **p** |
| μ | (Intercept) | -1.1 | 0.09 | -13 | 3.4e-36 |
| μ | StimulusCold | -0.26 | 0.094 | -2.7 | 0.0062 |
| μ | StimulusWarm | -0.38 | 0.099 | -3.8 | 0.00013 |
| μ | Burning_prob | 1.7 | 0.11 | 15 | 2.4e-47 |
| μ | Quartile | -0.03 | 0.074 | -0.41 | 0.69 |
| μ | Trial_n | -0.00079 | 0.00039 | -2 | 0.044 |
| μ | Stim_dur | 0.09 | 0.011 | 7.9 | 3.8e-15 |
| μ | StimulusCold:Burning_prob | -0.78 | 0.17 | -4.5 | 6.7e-06 |
| μ | StimulusWarm:Burning_prob | -0.47 | 0.18 | -2.5 | 0.011 |
| σ | (Intercept) | -0.21 | 0.093 | -2.3 | 0.023 |
| σ | StimulusCold | -0.09 | 0.098 | -0.92 | 0.36 |
| σ | StimulusWarm | -0.14 | 0.099 | -1.4 | 0.16 |
| σ | Burning_prob | -0.05 | 0.12 | -0.41 | 0.68 |
| σ | Quartile | 0.037 | 0.075 | 0.49 | 0.63 |
| σ | Trial_n | 0.00016 | 0.00039 | 0.42 | 0.68 |
| σ | Stim_dur | -0.017 | 0.012 | -1.4 | 0.15 |
| σ | StimulusCold:Burning_prob | 0.15 | 0.18 | 0.82 | 0.41 |
| σ | StimulusWarm:Burning_prob | 0.51 | 0.18 | 2.8 | 0.0058 |
| ν | (Intercept) | 1 | 0.53 | 1.9 | 0.059 |
| ν | StimulusCold | 0.76 | 0.49 | 1.6 | 0.12 |
| ν | StimulusWarm | 1.4 | 0.51 | 2.8 | 0.0054 |
| ν | Burning_prob | -7.4 | 0.86 | -8.6 | 7.7e-18 |
| ν | Quartile | -0.42 | 0.4 | -1 | 0.3 |

| VAS_Burn~Stimulus * Burning_prob + Quartile + Trial_n + Stim_dur + re(random = ~1 \| experiment/ID), ZOIB(link = logit) | | | | | |
|---|---|---|---|---|---|
| **parameter** | **contrast** | **β** | **SE** | **t** | **p** |
| ν | Trial_n | 0.0016 | 0.002 | 0.78 | 0.44 |
| ν | Stim_dur | -0.23 | 0.068 | -3.4 | 0.00071 |
| ν | StimulusCold:Burning_prob | 3.6 | 1.1 | 3.4 | 0.00065 |
| ν | StimulusWarm:Burning_prob | 3.8 | 1.1 | 3.5 | 0.00046 |
| τ | (Intercept) | -5.4 | 0.21 | -26 | 9.7e-136 |

Table 3, Burning ratings

*Thermosensory Index*

| coldwarm_ratio~Burning_prob + Quartile * Stimulus + Trial_n + Stim_dur + re(random = ~1 \| experiment/ID), ZOIB(link = logit) | | | | | |
|---|---|---|---|---|---|
| **parameter** | **contrast** | **β** | **SE** | **t** | **p** |
| μ | (Intercept) | 0.22 | 0.1 | 2.2 | 0.026 |
| μ | Burning_prob | -0.46 | 0.08 | -5.8 | 8.4e-09 |
| μ | Quartile | -0.89 | 0.11 | -7.7 | 1.3e-14 |
| μ | StimulusCold | 0.91 | 0.1 | 8.8 | 2.4e-18 |
| μ | StimulusWarm | -0.74 | 0.11 | -6.4 | 1.2e-10 |
| μ | Trial_n | -0.00076 | 0.00042 | -1.8 | 0.071 |
| μ | Stim_dur | 0.022 | 0.013 | 1.7 | 0.089 |
| μ | Quartile:StimulusCold | 0.47 | 0.19 | 2.5 | 0.014 |
| μ | Quartile:StimulusWarm | 0.48 | 0.2 | 2.4 | 0.018 |
| σ | (Intercept) | -0.24 | 0.1 | -2.3 | 0.023 |
| σ | Burning_prob | -0.29 | 0.089 | -3.3 | 0.0011 |
| σ | Quartile | -0.5 | 0.13 | -3.8 | 0.00014 |
| σ | StimulusCold | -0.21 | 0.11 | -1.9 | 0.06 |
| σ | StimulusWarm | 0.15 | 0.12 | 1.2 | 0.22 |
| σ | Trial_n | -0.00052 | 0.00045 | -1.2 | 0.25 |
| σ | Stim_dur | 0.056 | 0.013 | 4.4 | 1e-05 |
| σ | Quartile:StimulusCold | 0.92 | 0.2 | 4.6 | 5.5e-06 |
| σ | Quartile:StimulusWarm | 0.09 | 0.22 | 0.42 | 0.67 |
| ν | (Intercept) | 44 | 0.06 | 7.2e+02 | 0 |
| τ | (Intercept) | -1.5 | 0.082 | -18 | 2.5e-70 |

Table 4, Thermosensory index
